# MYT1L deficiency impairs excitatory neuron trajectory during cortical development

**DOI:** 10.1101/2024.03.06.583632

**Authors:** Allen Yen, Xuhua Chen, Dominic D. Skinner, Fatjon Leti, MariaLynn Crosby, Jessica Hoisington-Lopez, Yizhe Wu, Jiayang Chen, Robi D. Mitra, Joseph D. Dougherty

**Affiliations:** Department of Genetics, Washington University School of Medicine, Saint Louis, MO, USA; Department of Psychiatry, Washington University School of Medicine, Saint Louis, MO, USA; Edison Family Center for Genome Sciences and Systems Biology, Washington University School of Medicine, Saint Louis, MO, USA; Scale Biosciences, Inc. San Diego, CA, USA; DNA Sequencing and Innovation Lab, Washington University School of Medicine, Saint Louis, MO; Intellectual and Developmental Disabilities Research Center, Washington University School of Medicine, Saint Louis, MO, USA

**Author notes:** **Corresponding authors Current addresses: Lead Contact Information:** Dr. Joseph Dougherty Department of Genetics 4370 Duncan Ave. St. Louis, MO. 63110-1110.

## Abstract

Mutations that reduce the function of MYT1L, a neuron-specific transcription factor, are associated with a syndromic neurodevelopmental disorder. Furthermore, MYT1L is routinely used as a proneural factor in fibroblast-to-neuron transdifferentiation. MYT1L has been hypothesized to play a role in the trajectory of neuronal specification and subtype specific maturation, but this hypothesis has not been directly tested, nor is it clear which neuron types are most impacted by MYT1L loss. In this study, we profiled 313,335 nuclei from the forebrains of wild-type and MYT1L-deficient mice at two developmental stages: E14 at the peak of neurogenesis and P21, when neurogenesis is complete, to examine the role of MYT1L levels in the trajectory of neuronal development. We found that MYT1L deficiency significantly disrupted the relative proportion of cortical excitatory neurons at E14 and P21. Significant changes in gene expression were largely concentrated in excitatory neurons, suggesting that transcriptional effects of MYT1L deficiency are largely due to disruption of neuronal maturation programs. Most effects on gene expression were cell autonomous and persistent through development. In addition, while MYT1L can both activate and repress gene expression, the repressive effects were most sensitive to haploinsufficiency, and thus more likely mediate MYT1L syndrome. These findings illuminate the intricate role of MYT1L in orchestrating gene expression dynamics during neuronal development, providing insights into the molecular underpinnings of MYT1L syndrome.

## Introduction

Every brain cell shares the same genetic code, yet they exhibit a wide range of functions. This diversity arises because different cell lineages enact different gene expression programs that direct each cell in the embryonic brain to develop in a highly orchestrated manner. Disruption of these processes can lead to abnormal neurodevelopment and result in impaired cognition, communication, and adaptive behavior, as seen in profound autism and intellectual disability (ID)^1,2^. Notably, many genes associated with such neurodevelopmental disorders (NDDs) are expressed early during brain development and are involved in gene regulation and synaptic function^3,4^. Studies using post-mortem human brain tissue provide evidence that cortical excitatory neurons are commonly dysregulated in autism^5,6^. However, since these are end of life studies, whether this a cause or consequence of autism is unclear.

One such NDD associated gene is Myelin Transcription Factor 1 Like (MYT1L), which is highly expressed exclusively in postmitotic neurons in the embryonic brain and sustained at lower levels throughout life^7,8^. Early fibroblast-to-neuron transdifferentiation studies demonstrate that MYT1L promotes neuronal cell fate by repressing non-neuronal lineage programs^9,10^. Similarly, *in vivo* epigenetic studies of normal development show that MYT1L promotes neuronal differentiation by recruiting the SIN3B repressive complex to promoters and enhancers of postmitotic neurons to suppress early developmental programs^11^. Indeed, loss of MYT1L in multiple mouse models resulted in upregulation of a fetal gene expression signature^12–14^. To date, three pivotal studies have delved into the in vivo functions of MYT1L by creating transgenic mouse models. Each study uniquely disrupted a different exon of MYT1L (6 in Wohr *et al*.^15, 7^ in Chen *et al*.^12^, and 9 in Kim *et al*.^13^). The animal models are valuable tools to study the molecular and cellular consequences of MYT1L haploinsufficiency and the mice recapitulate many of the clinical presentations such as hyperactivity, structural malformations, obesity, and behavioral deficits^12–14^. However, it remains largely unknown how MYT1L haploinsufficiency influences the trajectory of neuronal differentiation *in vivo*, and whether the development of specific neuronal subtypes is particularly susceptible to the loss of MYT1L. Moreover, it is unclear if there is a critical moment in each cell’s developmental window during which MYT1L function is indispensable, as understanding this timeline could delineate when the transcriptional dynamics and developmental processes are amenable to interventions.

Detailed atlases mapping the gene expression profiles of thousands of cell types across the entire mouse brain have significantly advanced our understanding of brain organization under typical conditions^16–21^. Building upon this foundational knowledge, we can now explore how genetic perturbations affect neurodevelopment, specifically investigating the impact of disrupting a gene regulatory network through the loss of a single TF on this atlas. Given the widespread expression pattern of MYT1L in neurons, it is unclear if specific neuronal subtypes are more sensitive to MYT1L deficiency. Likewise, previous studies using bulk RNA sequencing have shown that MYT1L deficiency affects genes associated with the cell cycle^11,12,14^, differentiation^9,10^, and proliferation^22^. However, a limitation of bulk sequencing is that it only provides average gene expression data from a mixed population of cells, making it challenging to discern the precise origin of observed differences. For example, MYT1L haploinsufficiency results in an increased expression of developmental gene expression programs in vitro and in the post-natal brain^10,12,14^, but it remains unclear whether the observed differences are due to an increased proportion of immature progenitors or whether post-mitotic neurons are generated in proper numbers, but fail to mature completely and become trapped in an intermediate state. Furthermore, MYT1L functions as both a transcriptional repressor^10,23^ and activator^12,24^, but the variations in its role by cell type or developmental stage, as well as the sensitivity of the activated or repressed gene targets to disruption, are still unclear. Although loss of MYT1L leads to precocious differentiation during development^12^ and sustained activation of developmental programs in the adult brain^10,11^, the implications for neuronal development trajectory and cell-type specific fate specification remain unknown. Utilizing single cell transcriptomics, we can obtain a high-resolution mapping of dynamic developmental processes and elucidate how the loss of MYT1L contributes to the observed differential gene expression patterns.

In this study, we profiled a total of 313,335 nuclei to investigate the molecular and cellular consequences of MYT1L haploinsufficiency at the peak of neurogenesis (E14) and when neurogenesis is complete (P21). Our findings indicate that MYT1L deficiency primarily impacts excitatory neurons. We further identified that genes regulated by MYT1L, whether activated or repressed, exhibit cell type-specific responses to MYT1L haploinsufficiency. A significant number of dysregulated genes were TFs or epigenetic regulators temporally expressed during specific time windows, highlighting lineage specific gene regulatory networks. In summary, our findings provide insights on how MYT1L haploinsufficiency disrupts embryonic and postnatal neurodevelopment. We have identified key transcriptional networks and defined the vulnerable cell types and developmental stages that potentially contribute to the pathogenesis of MYT1L syndrome.

## Results

### Loss of MYT1L disrupts proportions of excitatory and inhibitory neurons

To characterize the role of MYT1L during peak neurogenesis and to understand the acute consequences of MYT1L haploinsufficiency and loss on cell fate specification and maturation, we applied a combinatorial indexing approach^25,26^ to profile and analyze transcription from 216,830 nuclei from the developing forebrain of embryonic day 14 (E14) MYT1L knockout (KO), heterozygous (Het), and wild type (WT) animals (**Figure 1A, B**). We find that all cell types are well represented across all genotypes (median genotype LISI score^27^=2.7) (**Figure 1C**). We identified 26 clusters representing 7 broad neural cell types, which were further classified into three subtypes of radial glial cells (*Hes1* and *Nestin* positive), 3 subtypes of intermediate progenitor cells (*Neurog2* and *Eomes* positive) fated to be excitatory neurons, 3 subtypes of inhibitory intermediate progenitor cells (*Dlx1* and *Nkx2.1* positive), 8 subtypes of excitatory neurons (*Neurod6* and *Tbr1* positive), 9 subtypes of inhibitory neurons (*Gad1* and *Gad2* positive), Cajal-Retzius cells, oligodendrocyte progenitor cells, and microglia (**Figure 1C, D**). We assigned cell cycle scores based on cell cycle phase marker gene expression and confirmed that the progenitors were mostly in G2M or S, while the post-mitotic neurons were in G1/G0 (**Figure 1E**) and expressed MYT1L (**Figure 1F**). The progenitor cells segregated into two distinct populations—the root clusters which gave rise to divergent excitatory and inhibitory neuron developmental trajectories. This profile of cellular diversity indicated that we captured a developmental window encompassing differentiation and maturation processes, enabling us to investigate the molecular and cellular consequences of loss of MYT1L in the developing E14 cortex.

**Figure 1.**
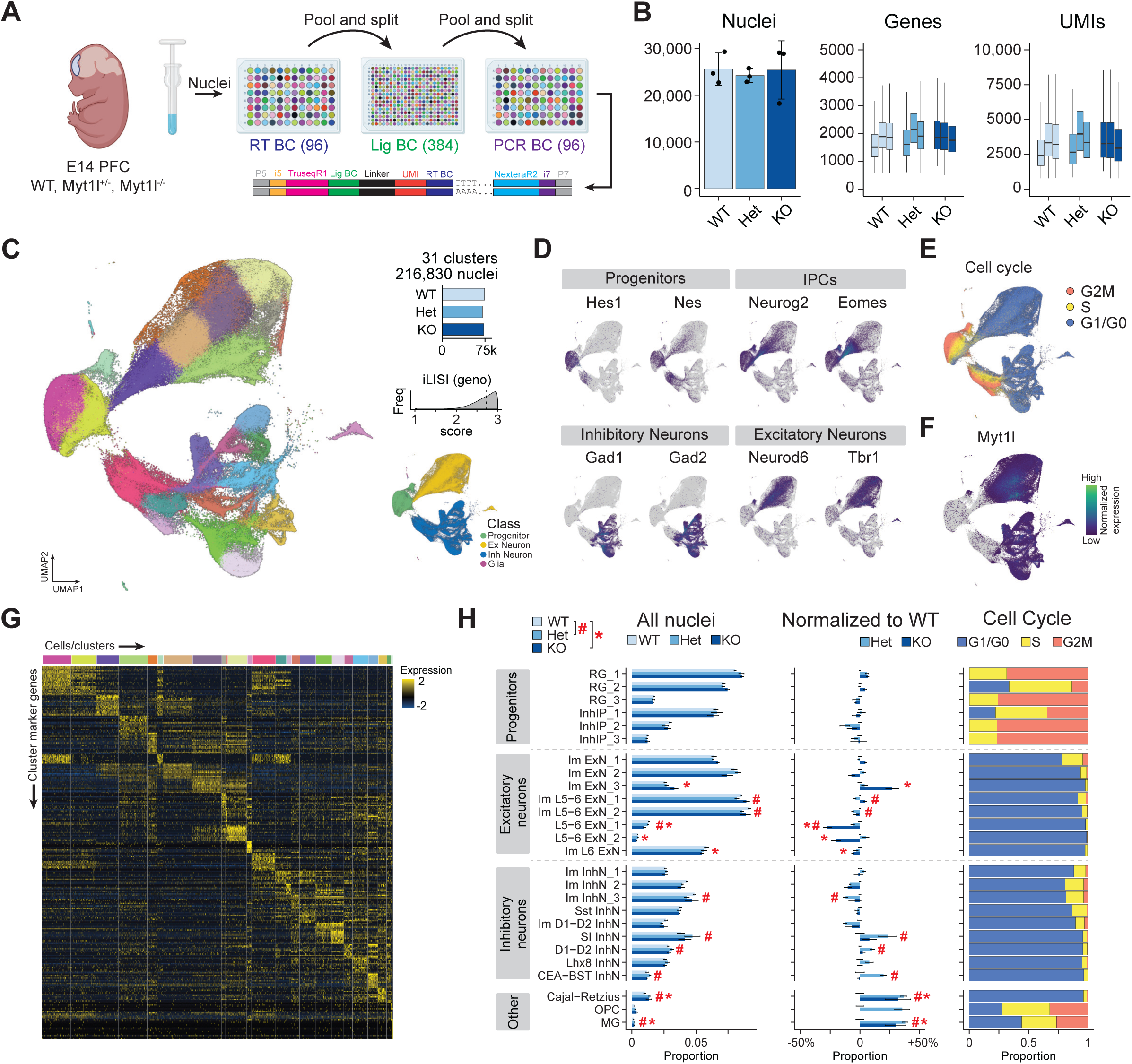
Single nucleus transcriptional profiling of E14 forebrain in MYT1L animals. (A) Schematic showing dissection of forebrain tissue, isolation of nuclei, combinatorial barcoding, and generation of snRNAseq libraries. **(B)** General library statistics showing average nuclei per genotype, median genes per nuclei, and median UMIs per nuclei. **(C)** Uniform manifold approximation and projection (UMAP) of 216,830 nuclei from MYT1L WT, Het, and KO animals colored by cell type. The bar plot shows the total number of nuclei per genotype across biological replicates. The histogram shows the local inverse Simpson’s index (LISI) score has a median of 2.7, indicating that the genotypes are well mixed and integrated. The bottom right UMAP inset shows all nuclei color coded by cell class. **(D)** Top markers for progenitors, intermediate progenitor cells (IPCs), inhibitory neurons, and excitatory neurons. **(E)** UMAP of all nuclei color coded by cell cycle score based on cell cycle genes. **(F)** UMAP plot showing expression of MYT1L in postmitotic excitatory and inhibitory neurons. **(G)** Heatmap showing the top marker gene expression (rows) for cells in each cluster (columns). **(H)** Bar plots show the average relative proportions of nuclei in each annotated cell cluster for MYT1L WT, Het, and KO genotypes (left). These proportions are normalized to WT (center). The composition of each cluster by cell cycle phase is shown on the right.

Because MYT1L is highly expressed in virtually all neurons during neurogenesis (**Figure 1F**), we sought to assess the short-term consequences of its deficiency on overall cell type proportions. We found subtle but statistically significant disruptions to the abundance of post-mitotic immature excitatory neurons (Im ExN_3), deep layer excitatory neurons (Im L5-6 ExN_1, Im L5-6 ExN_2, L5-6 ExN_1, L5-6 ExN_2, and Im L6 ExN), immature inhibitory neurons (Im InhN_3), and specific subtypes of inhibitory neurons (somatosensory cortex (SI), Darpp32+ D1-D2, and CEA-BST) (**Figure 1H**). Radial glia and inhibitory intermediate progenitors were mostly unaffected by the loss of MYT1L. Non-cycling immature excitatory neurons (Im ExN_3) in the subventricular zone (SVZ) were the most developmentally immature cells from the excitatory trajectory that were affected, showing an increase in abundance in KOs compared to WT which could be a result of precocious neuronal differentiation.

### Loss of MYT1L disrupts excitatory neuron development

We then conducted a differential expression analysis to analyze the molecular signatures of each cell type and determine which subtype exhibited the most significant transcriptional changes due to MYT1L deficiency. Individual clusters were aggregated into pseudobulk groups and then we used DESeq2^28^ to conduct pairwise analyses between WT and KO genotypes to uncover the most pronounced expression differences within each cell type. We identified 1,174 unique differentially expressed genes (DEGs; BH adjusted P-value < 0.1; expression level change ≥ 15%) between WT and KO, of which 781 were upregulated in KO cells and 415 were downregulated compared to WT (**Figure 2A**). There were only 11 genes that were not exclusively up or downregulated across all cell types. DEGs that were found to be unique to a single cluster accounted for 54% (633/1174) of the DEGs, demonstrating disruption of both ubiquitously expressed and cell type-specific genes. Notably, deep layer excitatory neurons, especially immature L6 neurons, harbored the majority of DEGs (**Figure 2A**), indicating that these neurons may be particularly sensitive to loss of MYT1L, which is consistent with their disrupted cell proportions (**Figure 1H**). Progenitor cells, which do not yet express MYT1L, showed no DEGs, demonstrating that the molecular and cellular consequences of MYT1L deficiency are cell intrinsic to the cells that express MYT1L. This suggests that there are no signals from the differentiating neurons that robustly influence the transcriptional identity of the proliferating progenitor pool at E14.

**Figure 2.**
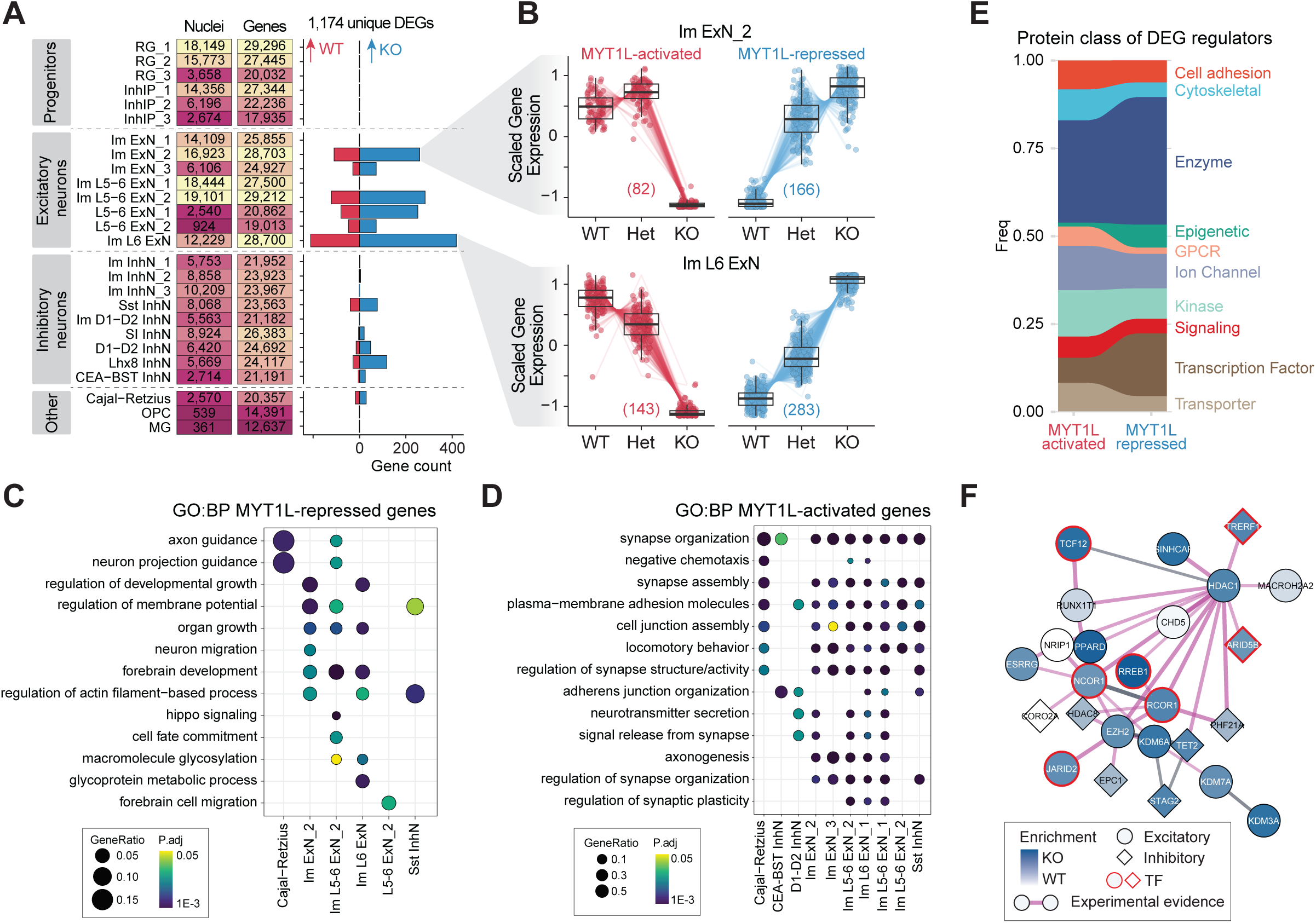
Cell type-specific changes in gene expression. (A) Summary plot showing the numbers of nuclei and genes detected in each cluster. The bar plot shows the number of differentially expressed genes that are upregulated in WT (red) or upregulated in KO (blue). **(B)** MYT1L gene dose-dependent gene expression patterns of DEGs in Im ExN_2 and Im L6 ExN clusters separated by those that are MYT1L-activated (loss of expression in KO) and MYT1-represssed (gain of expression in KO). **(C, D)** Dotplot showing enriched GO biological processes in MYT1L-repressed **(C)** and MYT1L-activated **(D)** DEGs. **(E)** Plot showing the frequency of annotated protein classes of the DEGs. **(F)** STRING physical protein-protein interaction networks for a coregulatory module. Darker blues indicate the gene was higher expressed in KO, a red outline signifies that the gene is a TF, and the shape indicates the cell type.

Given that MYT1L homozygotes do not survive postnatally, and the human disorder is caused by haploinsufficiency, it is of interest to analyze Hets. Consequently, we investigated whether the DEGs were dose-responsive to the number of MYT1L copies, or if there were non-linear effects of MYT1L loss. Furthermore, whether this pattern of regulation was the same for activated and repressed genes may suggest which function is most critical to the disorder. Therefore, we modeled the number of functional alleles as an ordinal factor and classified 522 genes that were upregulated in KOs as MYT1L-repressed genes, while the 451 genes that were upregulated in WTs were considered MYT1L-activated genes. We found that MYT1L-repressed genes were more sensitive to the gene dose of MYT1L than genes activated by MYT1L (**Figure 2B**). In MYT1L-activated genes, expression levels in Hets were similar to WTs suggesting that these target genes are sufficiently activated even with decreased levels of MYT1L. However, the MYT1L-repressed genes exhibited a nearly linear gene-dosage response and in some cases Hets were more similar to KOs (**Figure 2B**). This suggests that MYT1L repressed targets become highly upregulated with the loss of a single MYT1L allele.

Prior bulk RNAseq of the E14 MYT1L Het mouse cortex revealed an immature transcriptional signature when compared to WT^12^. To evaluate if a particular cell type was driving this effect, we performed gene ontology (GO) enrichment analysis for the DEGs in each cluster. We found that MYT1L-repressed genes (KO>WT) are represented in development, neuron migration, and cell fate commitment pathways and were the top enriched pathways in MYT1L KO neurons (**Figure 2C**), while MYT1L-activated genes (WT>KO) are involved in synapse organization, axonogenesis, and neurotransmitter secretion and transport (**Figure 2D**). Together, this revealed that loss of MYT1L results in an immature developmental transcriptional state. By analyzing the DEGs across cell types, we found that MYT1L-repressed genes had a more functionally diverse response to loss of MYT1L compared to those that are activated by MYT1L. This suggests that the suppression of developmental genes is critical to ensure proper neuronal maturation.

We next asked if our lists of MYT1L-activated and -repressed genes are direct or indirect targets of MYT1L. To test this, we integrated our E14 snRNA-seq dataset with an age and region-matched E14 forebrain MYT1L CUT&RUN dataset that cataloged 560 high-confidence MYT1L binding sites within promoter sequences^11^. 58 direct MYT1L targets were found to be differentially expressed and most of these showed dose-dependent responses to MYT1L. Comparing KOs to WTs, 47 (36 were dose-dependent) were upregulated and 16 (9 were dose-dependent) were downregulated consistently across cell types, reinforcing that MYT1L functions as a transcriptional repressor at ∼80% of consistent targets. Additionally, this demonstrates that the DEGs were largely driven by indirect effects of MYT1L deficiency. Indeed, differentially expressed MYT1L targets were significantly enriched for TFs (81/560; P=2.2x10^-16^, Fisher’s exact test) (**Figure 2E**).

We then performed network analysis on all the DEGs identified through cluster pseudobulk analysis to gain insight into functional interactions and putative upstream regulators^29^. We identified modules by clustering the protein-protein interaction network based on functional annotations and found that the modules were significantly enriched with transcription regulators and epigenetic factors. Many of these had higher expression in the L5-6 ExN_1 cluster from MYT1L KO animals (**Figure 2F**). This provides evidence that MYT1L can be a transcriptional regulator that not only influences its direct target genes, but also downstream indirect targets within a gene network.

As studies have shown that some autism risk genes disrupt excitatory and inhibitory neurogenesis, we then extended our analysis to test if our observed transcriptional disruptions converge with genes associated with neurodevelopmental disorders. We intersected the excitatory and inhibitory neuron DEGs (**Figure 2A**) with a list of 932 high-confidence autism-related genes from the SFARI database with a score of 1 or 2 ^30–32^. We observed a significant overlap between DEGs and the SFARI genes, with 178 out of 1174 DEGs (15.2%) being shared (P=2.1x10^-12^, chi-square test with Yates’ continuity correction). This overlap comprised 146 genes from the excitatory neuron clusters and 32 genes were from the inhibitory neuron clusters. As autism genes have a bias towards genes highly expressed in neurons^33^, we wanted to test if this overlap was merely driven by a neuron bias. Therefore, we randomly sampled 932 highly expressed genes in these clusters a thousand times and examined the overlap with SFARI genes. The median overlap was 14 compared to the 178 seen here, indicating a greater than 10-fold enrichment for SFARI genes among MTY1L DEGs. Deeply examining the 178 genes, many of the autism associated DEGs such as *Ext1* and *Phf21a* exhibited pronounced effects in deep layer excitatory neurons (**Supplemental Figure 1**). This finding indicates that the pathways perturbed by MYT1L deficiency have a signature similar to pathways disrupted by a subset of autism genes involved in axon guidance, neuronal migration, and chemical synaptic transmission, suggesting convergence with key autism-related genes and pathways^6^. Overall, these observed transcriptomic changes reveal molecular changes that preferentially affect the maturation and function of deep layer excitatory neurons.

### Loss of MYT1L disrupts transcriptional maturation

Our differential analysis shows that MYT1L deficiency is associated with an immature transcriptional signature in excitatory neurons. While an explanation is that these genes are simply dysregulated, we hypothesize that MYT1L deficient neurons progress along their developmental trajectories at a slower pace, resulting in an immature gene signature. Additionally, we hypothesize that there is a critical moment during differentiation when MYT1L function is most needed to guide the developmental trajectory. To test these hypotheses, we assessed the differences in maturation trajectories leading up to and through this developmental window between genotypes. We used Monocle3^34^ to reconstruct a pseudotemporal trajectory (**Figure 3A**) independent of our prior cluster definitions. This models the cell state as a continuum of dynamic changes, allowing us to quantify gene expression changes as the cell progresses through differentiation. We observed subtle yet widespread disruption to the distribution of Het and KO nuclei compared to WT in pseudotime states (**Figure 3B-D**). This suggests a developmentally immature signature that can be missed when looking only at cell proportion based on cluster markers.

**Figure 3.**
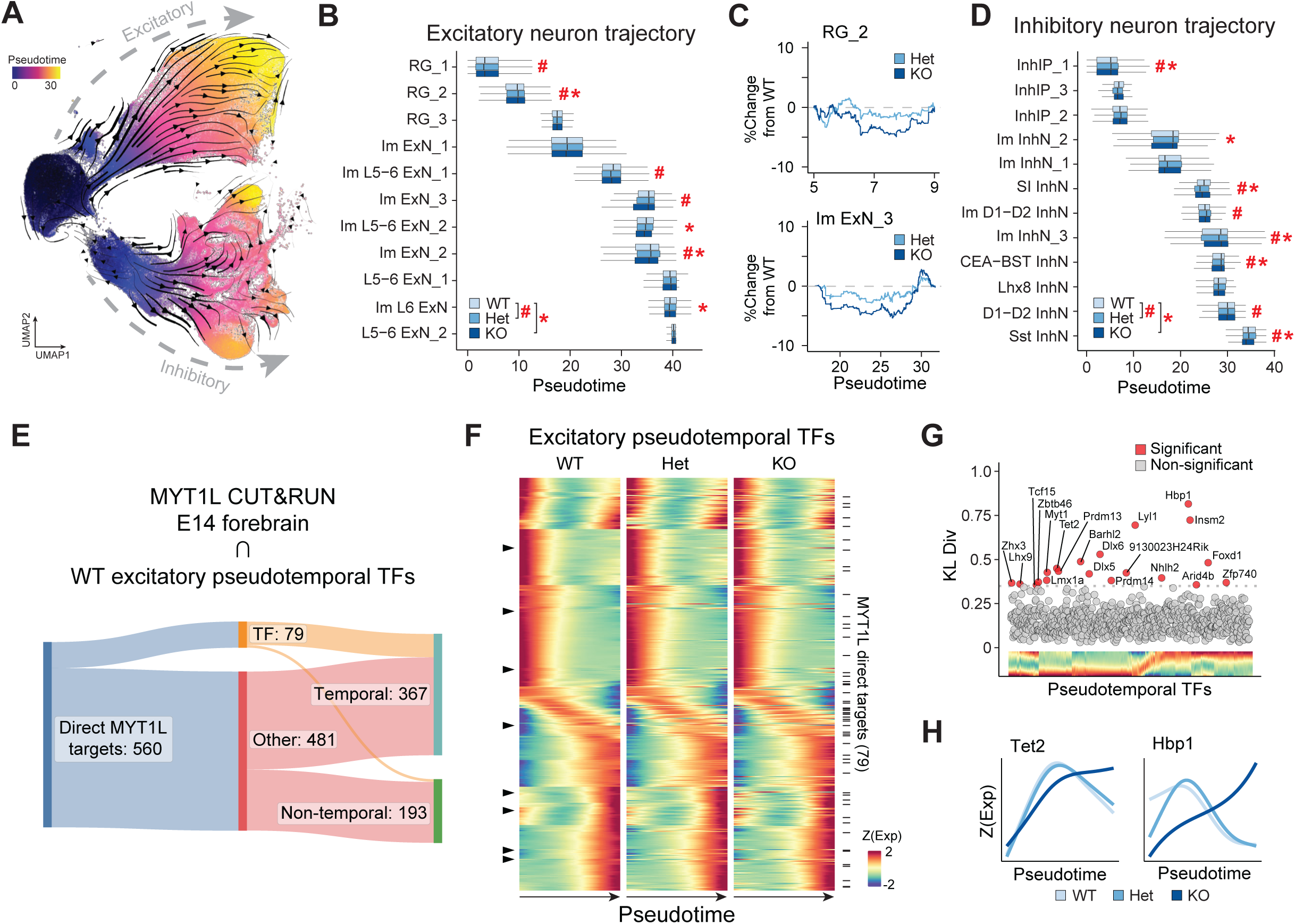
Loss of MYT1L disrupts excitatory neuron maturation. (A) UMAP plot of all nuclei from all genotypes colored by pseudotime and overlaid with RNA velocity trajectories. **(B)** Boxplots showing distributions nuclei from excitatory neurons along pseudotime. Kolmogorov-smirnov tests were performed to test for differences in distributions. P<0.05 for WT and Het comparisons are noted with a #, while statistically significant WT and KO comparisons are noted with a *. **(C)** Representative plots showing the relative differences of distribution in the RG_2 and Im ExN_3 clusters of MYT1L Het and KO nuclei compared to the WT distribution. **(D)** Boxplots showing the distributions of inhibitory neuron nuclei along pseudotime. P<0.05 for WT and Het comparisons are noted with a #, while statistically significant WT and KO comparisons are marked with a *. **(E)** Diagram showing the number of direct MYT1L targets identified by CUT&RUN that had a dynamic gene expression profile across pseudotime. **(F)** Heatmaps showing scaled expression of WT (left), Het (middle), and KO (right) excitatory neuron pseudotemporal genes. Each row represents a gene and sorted according to their expression peak in pseudotime. The black tick marks on the right note the rows in which genes are MYT1L direct targets determined by CUT&RUN in **E**. Black triangles on the left denote a subset of genes as examples with disrupted timing of expression. **(G)** Scatterplot showing the Kullback-Liebler divergence metric to identify differential pseudotemporal expression profiles in KOs compared to WT. **(H)** Representative traces of the differential pseudotemporal gene expression profiles for Tet2 and Hbp1 across genotypes.

To identify drivers of excitatory neuron development, we analyzed the expression of TFs along pseudotime in WT cells, providing us with a putative timeline of gene activation and expression from progenitors to differentiated excitatory neurons (**Figure 3E**). Using these temporal profiles, we can then test if loss of MYT1L causes variations in the timing of expression of specific TFs which could profoundly impact the developmental trajectory of excitatory neurons. We found 27 TFs that showed disrupted timing of expression as a result of MYT1L deficiency using the Kullback-Leibler divergence test (**Figure 3F,G**). These TFs were generally de-repressed in Hets and KOs and are involved in developmental regulation (*Dlx5*, *Dlx6*, and *Hoxd10*), control of cell cycle progression (*Hbp1*), neurogenesis (*Nhlh2*, *Lmx1a*, and *Insm2*), and epigenetic regulation (*Tet2* and *Prdm*). To identify where MYT1L may have the greatest effect, we intersected all the excitatory pseudotemporal TFs with the E14 MYT1L CUT&RUN peaks^11^ and found an enrichment of direct MYT1L targets during a transient period shortly after the transition from progenitor to postmitotic neuron, suggesting its important role during this critical moment. We also found that six genes within this pseudotime bin (*Efna4, Ccng2, Nbr1, Frmd4b, Sorsb2*, and *Midn*) were targets of ZBTB12, a molecular gatekeeper known to safeguard the unidirectional transition of progenitors to differentiated states^35^. Together, this provides a pseudotime-resolved sequence of MYT1L target gene expression, and identification of a critical developmental window, where alterations in these patterns during this sensitive moment may lead to disruptions in neuronal differentiation and maturation.

### Sensitivity of excitatory neurons persist throughout neurodevelopment

To investigate the long-term effects of MYT1L deficiency on both cell proportion and transcriptional changes, we conducted a snRNAseq analysis of the cortex in juvenile male and female WT and MYT1L Het animals at P21, when neurogenesis is largely completed. Analysis of KOs are not possible as they are not viable postnatally^12–14^. We analyzed snRNAseq data from 96,505 nuclei to identify 19 types of excitatory neurons spanning cortical layers, 11 subtypes of inhibitory neurons, and 8 non-neuronal types using hierarchical correlation mapping, referencing the taxonomies and subclass annotations from the Allen Brain Cell (ABC) Atlas^19^ (**Figure 4A**, **Supplemental Figure 2**). When analyzing overall proportions of excitatory neurons, we observed significantly fewer L2/3 IT ENT and L4/5 IT neurons in MYT1L Het cortices compared to WT, while there were increased numbers in L6 CT, L6 IT, and L6b/CT neurons in the Hets (**Figure 4B**). By P21, 985 cluster pseudobulk DEGs were detected, of which 576 were unique to a single cell type and nearly exclusive to excitatory neurons (**Figure 4C**, **Supplemental Figure 3**). Similar to the E14 DEGs, we observed an increased number of upregulated DEGs upon loss of MYT1L in Hets, indicating de-repression. L6 neurons were the most affected, with modest effects on upper L2/3 intrathalamic (IT) cortex and mid-layer L4/5 IT neurons. 48% (280/576) of these DEGs overlapped with MYT1L CUT&RUN targets from adult prefrontal cortex^11^ (**Figure 4C**). This significant overlap implies that a substantial portion of the DEGs observed at P21 may be directly influenced by MYT1L binding to their promoter regions. GO analysis revealed that DEGs upregulated in WT were associated with axon guidance, synaptic cell adhesion, and neurotransmission (**Figure 4D**), while genes upregulated in Hets were enriched in pathways related to nervous system development and axonogenesis (**Figure 4E**). This reflects the E14 enrichment analysis and demonstrates that MYT1L Het excitatory neurons are immature compared to WT and that while the magnitude of effect on neurodevelopment and maturation is greatest during embryonic development, some deficiency is sustained throughout early postnatal development.

**Figure 4.**
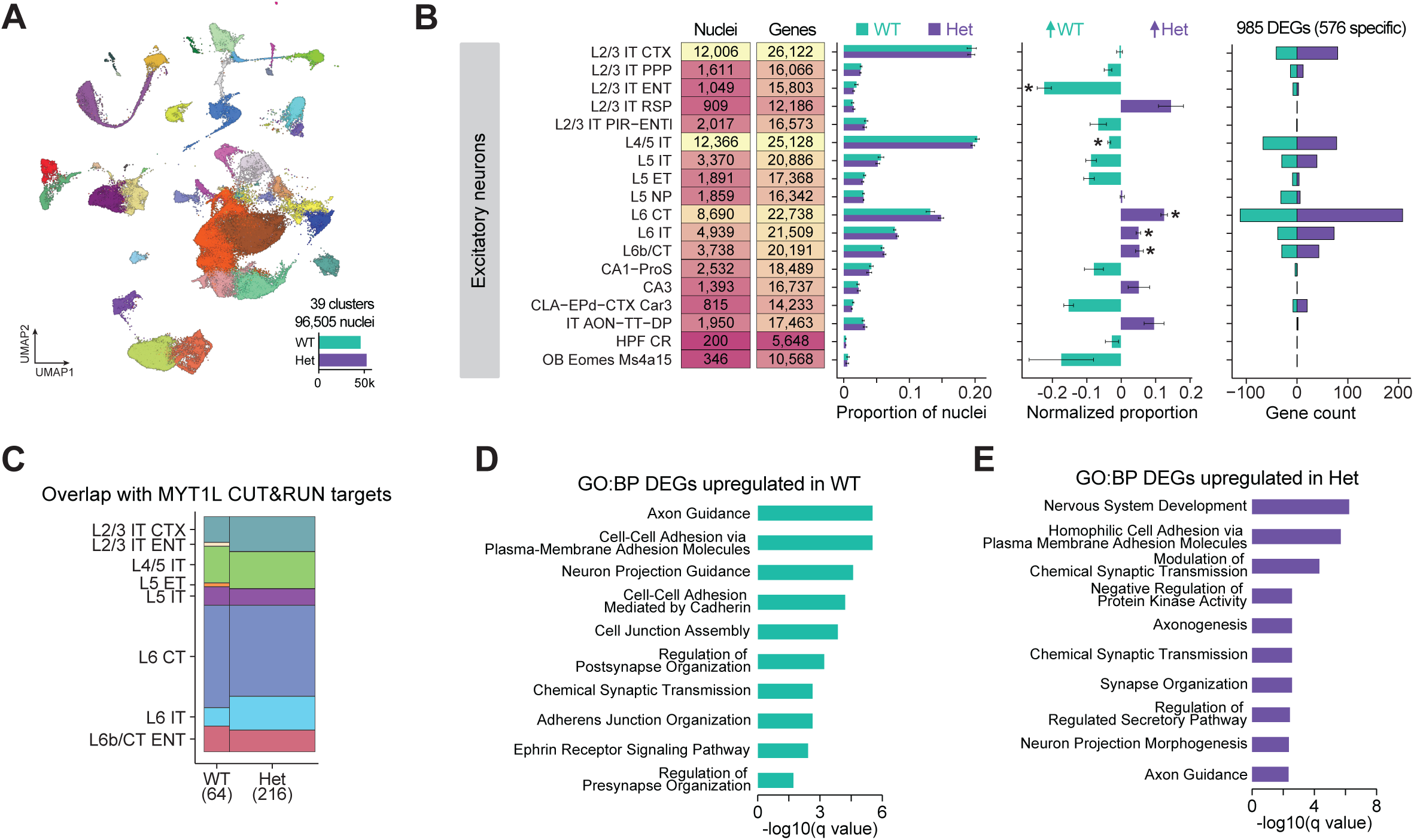
Single nucleus transcriptional profiling of P21 cortex in MYT1L animals. (A) UMAP projection showing 96,505 nuclei in 39 clusters from MYT1L WT and Het animals. **(B)** Summary plot showing the numbers of nuclei and genes detected in each cluster (left). Bar plots show the average relative proportions of nuclei in each annotated excitatory neuron cluster for MYT1L WT and Het genotypes. These proportions are normalized to WT (center bar plot). The right bar plot shows the number of pseudobulk differentially expressed genes that are upregulated in WT (cyan) or upregulated in Het (purple). **(C)** Mosaic plot showing the proportions of DEGs that overlap with MYT1L direct targets identified by CUT&RUN. The total numbers of overlapping genes are indicated in parentheses below the genotype labels. **(D, E)** GO analysis of biological processes of DEGs that are upregulated in WT and upregulated in Hets.

To deepen our understanding of the developmental progression from E14 progenitors to terminally differentiated cell types at P21, we integrated the two datasets together, analyzing a total of 313,335 nuclei. This integration revealed distinct developmental pathways for excitatory and inhibitory neurons, branching out from the clusters of progenitors (**Figure 5A**). For the subsequent analyses, we focused on the excitatory neuron trajectory encompassing 191,217 nuclei. We found that the E14 L5-6 ExN_1 and Im L6 ExN clusters were transcriptionally similar to the P21 L6 and L6b corticothalamic (CT) clusters and were observed at a transition zone between the ages. This indicates that the deep layer neurons are the first to exhibit markers indicative of a terminally differentiated cell type. The E14 Im ExN clusters showed a developmental trajectory towards the upper layer cortical neurons.

**Figure 5.**
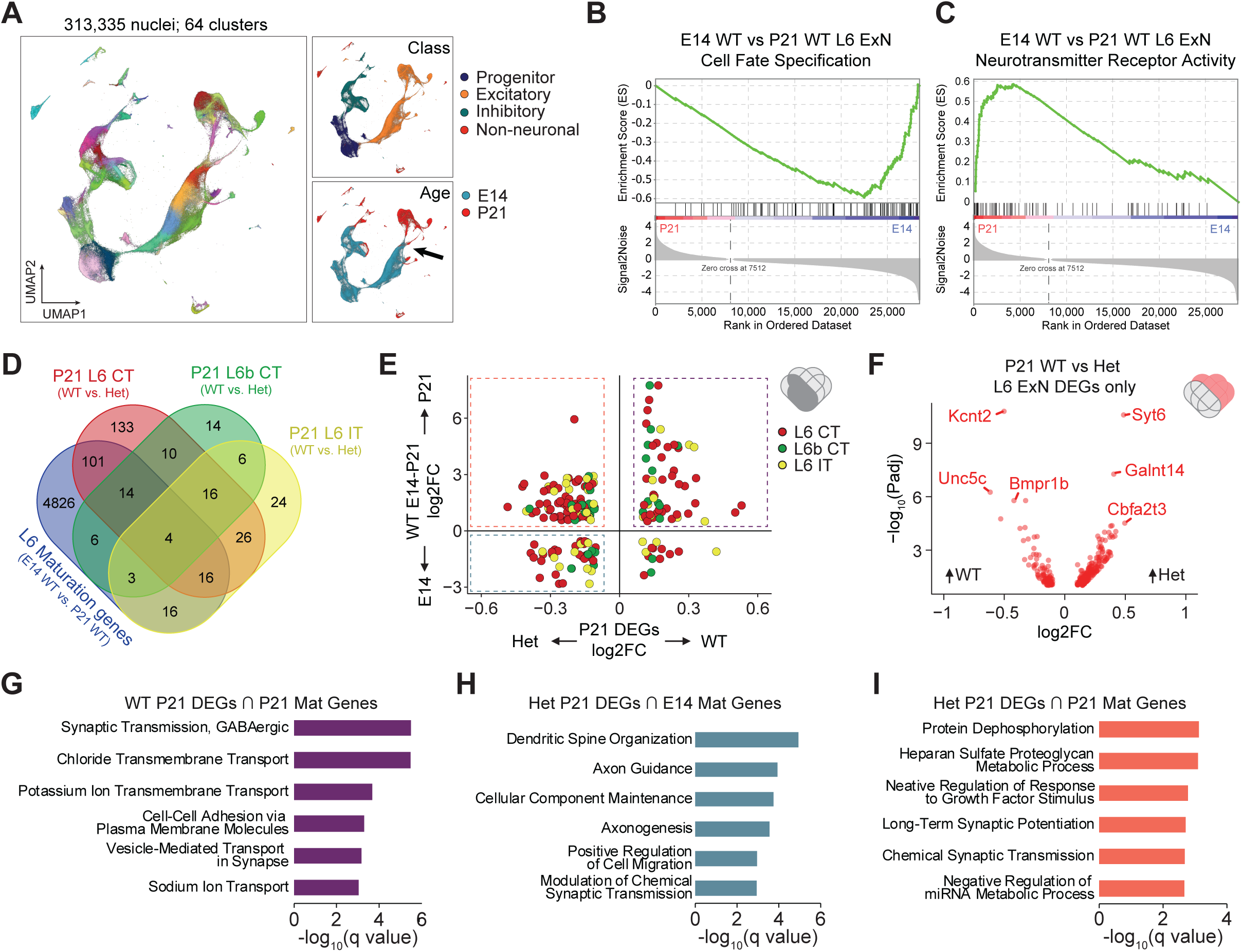
Integrated analysis of E14 and P21 nuclei. (A) UMAP projection showing integrated data from E14 and P21 datasets. The top right inset shows the cells colored by cell class. The bottom right inset shows the cells colored by age, with an arrow indicating the L6 transition zone. **(B,C)** GSEA analysis of the expression dataset comparing E14 L6 ExN to P21 L6 ExNs. In **B**, the green line shows the de-enrichment of genes in the Cell Fate Specification GSEA list in P21 L6 ExNs compared to E14 L6 ExNs. In **C**, the plot shows enrichment of genes associated with Neurotransmitter Receptor Activity in P21 compared to E14. **(D)** Venn diagram showing the number of overlapping genes in the maturation-associated gene set (E14 WT L6 ExN vs. P21 WT L6 ExNs) with DEGs from WT and Het P21 L6 clusters. **(E)** Scatterplot comparing the magnitude and direction of effect of the common genes between the WT L6 maturation genes and P21 L6 ExNs (shaded region indicated in venn diagram cartoon in the upper right corner). The x axis represents the log2FC from WT and Het P21 DEGs from the 3 L6 ExN clusters. The y axis is the log2FC of WT E14 L6 ExN and WT P21 L6 ExN DEGs. The upper right quadrant (purple) are genes upregulated in P21 WT and also generally at P21. The upper left (orange) quadrant are genes upregulated in P21 Hets that are also generally increased at P21. The bottom right (teal) are genes that are upregulated in P21 Hets, that are upregulated at E14. **(F)** Volcano plot showing the log2 fold change and direction of effect of DEGs from P21 L6 ExN DEGs that are not shared with the E14-P21 WT maturation gene list. **(G-I)** GO analyses of DEGs that are in the selected quadrants of **E**.

We next sought to test the hypothesis that MYT1L heterozygosity disrupted cell-type specific transcriptional maturation by P21. First, to understand the biological processes underlying the WT maturation of immature E14 L6 ExNs compared to their mature counterparts at P21, we performed gene set enrichment analyses (GSEA) on their gene expression profiles. As expected, we observed a de-enrichment of cell fate specification genes and an enrichment of neurotransmitter receptor activity genes in P21 L6 neurons (**Figure 5B,C**). Next, through differential gene expression analysis, we identified 4986 genes with significant differences between WT E14 and WT P21 L6 ExNs, highlighting a signature for neuronal maturation. Then, we further investigated the impact of MYT1L deficiency on the expression of these maturation-associated genes by comparing these 4986 genes with DEGs in P21 Het and WT L6 ExNs. Our analysis revealed that 35-42% of the P21 DEGs in the MYT1L deficient neurons overlapped with the maturation gene set (**Figure 5D**) (P=1.3x10^-4^, chi-square test with Yates’ continuity correction), indicating that about half of the transcriptional effects of MYT1L deficiency can be summarized as a disruption in neuronal maturation programs. To get more insight into the disrupted pathways, we analyzed these shared maturation genes (**Figure 5E**) as well as the genes that were only dysregulated at P21 (**Figure 5F**). We found that P21 DEGs upregulated in WTs showed an overall increase of expression of maturation-associated genes at P21 (upper right quadrant) and are related to synaptic transmission, GABA signaling, and ion transport (**Figure 5G**), while only a few E14 L6 maturation genes were upregulated (**Figure 5E**, lower right quadrant). In contrast, there was a significant number of P21 DEGs upregulated in Hets that showed higher expression of maturation genes at E14 (**Figure 5E**, lower left quadrant) which were related to synapse organization, axon guidance, and regulation of cell migration (**Figure 5H**). Finally, we observed a significant number of P21 Het DEGs that showed higher expression at P21 (**Figure 5E**, upper left quadrant), suggesting that MYT1L deficiency results in atypical expression of some developmental gene programs. GO analysis revealed an enrichment in pathways related to protein dephosphorylation and proteoglycan processes. Moreover, genes associated with semaphorin receptor binding genes, specifically *Sema4a*, *Sema7a*, and *Sema4d* were found to be upregulated. Notably, *Sema4d* is recognized for its role as an intrinsic inhibitor of axonal pathfinding^36–38^. This suggests that elevated levels of *Sema4d* in MYT1L Hets may impair axonal development and formation of synaptic connections. Collectively, this integrated analysis demonstrates that the P21 Het L6 ExNs exhibit an immature transcriptional signature, and the dysregulated genes suggests disrupted axon development and neurotransmitter signaling. The convergence of these findings underscores the critical role of MYT1L in guiding neuronal maturation and establishes a link between MYT1L heterozygosity and the perturbation of essential developmental pathways in L6 excitatory neurons.

## Discussion

In this study, we analyzed the transcriptomes of 313,335 nuclei across neurodevelopment in a model of MYT1L mutation, allowing us to delineate the molecular and cellular consequences of loss of MYT1L. Leveraging single-cell atlases of the mouse brain as references, we have advanced our understanding of how the disruption of a single TF can perturb neurodevelopment and maturation processes. We implemented pseudotime and RNA velocity to quantitatively assess neuronal transcriptional maturation. Our results reveal that although MYT1L is expressed in all neurons, deep layer excitatory neurons are particularly susceptible to MYT1L haploinsufficiency, resulting in an immature transcriptomic signature. This signature can be a result of precocious differentiation of earlier progenitors, a slower transition from progenitors into excitatory neurons, or cells stalled in a partially differentiated state. This deficiency causes a delay in neuronal maturation at E14, and the dysregulation of the regulatory programs that control neuronal maturation persist through P21. We also demonstrate that MYT1L primarily functions as a transcriptional repressor, affecting gene expression programs linked to key developmental processes like axon guidance, neuron migration, and cell fate commitment. We found these MYT1L-repressed pathways were gene dose-responsive, with even slight reductions in MYT1L levels leading to substantial upregulation of these genes. In contrast, genes activated by MYT1L, mainly those involved in synaptic function and neurotransmission, were more tolerant of haploinsufficiency. Additionally, our findings show that the dysregulated genes were enriched with TF and epigenetic regulators, which can initiate a cascade of downstream effects stemming from MYT1L perturbation. Using a brain region and age matched MYT1L CUT&RUN dataset, most effects at E14 were indirect and the percentage of direct effects increased at P21. Because MYT1L recruits the SIN3B deacetylation complex, it is possible that many of the “indirect” regulatory targets as measured by CUT&RUN are in fact direct as the deacetylated histones can have repressive epigenetic “memories”^39^, and a deeper analysis of histone state may disentangle this. Nonetheless, the overall results highlight the cell type-specific and developmental stage-specific nature of MYT1L’s function.

To date, there are three transgenic mouse models that disrupt different exons of MYT1L that converge on a hyperactivity phenotype, while other behaviors are varied likely because different assays were used^12,13,15^. A study by Weigel et al.^14^ performed scRNAseq on the neonatal (P0) forebrain from the mice described in Wohr et al.^15^ and found a decreased number of newly formed neurons in the subventricular zone. Additionally, the authors observed an increased expression of non-neuronal gene expression programs which can perturb neuronal cell identity. Here, we observed a slight upregulation of mouse embryonic fibroblast (MEF) signature genes at P21, albeit with a minor effect size. They also show L5/6 neurons as having the greatest number of DEGs, consistent with the work here. Interestingly, they also observed a moderate increase in the proportion of striatal inhibitory neurons in MYT1L Hets, however, additional experiments are needed to interpret these findings.

In the current era of genomics bulk RNAseq and snRNAseq are primary techniques for examining transcriptional landscapes across various biological perturbations. However, the concordance between DEGs identified through bulk and single-cell approaches is generally low. This discrepancy can be attributed in part to factors such as RNA capture efficiency, gene dropouts, and data sparsity. By leveraging a previously reported bulk RNAseq dataset from Chen et al.^12^, we perform a pairwise comparison with the pseudobulk aggregated snRNAseq dataset to discern whether the noted differences stem from variations in cell proportions or from intrinsic transcriptional changes within each cell type. In analyzing our E14 data, we observe that the discrepancies in cell proportions for MYT1L Het samples compared to WT are generally within a 10% margin, albeit with some exceptions, including the Im ExN_3, L5-6 ExN_1, and L5-6 ExN_2 clusters. Upon examining clusters that exhibit a substantial number of pseudobulk DEGs, such as Im L6 ExN, Im ExN_2, and Im L5-6 ExN_2, we identify only minor shifts in cell proportions. For these specific clusters, it appears that differential expression is predominantly driven by transcriptional alterations within the cell types, rather than changes in their proportions.

While MYT1L is a neuron-specific TF, it remains uncertain whether its loss may exert non-cell autonomous effects on surrounding glia during postnatal development. Our analysis revealed a relatively higher proportion of oligodendrocytes and microglia in P21 MYT1L Hets compared to WT, but we did not detect any DEGs in these cell types. However, with relatively low numbers of cells and gene counts in these clusters, there may be differences below our threshold to detect. These findings suggest the possibility of MYT1L-mediated effects on oligodendrocytes and microglia, yet further investigations with increased cell numbers are needed to elucidate the nature and extent of these effects. What was abundantly clear in our data at both time points was the profound, cell type-specific transcriptional responses to MYT1L deficiency, especially in deep layer excitatory neurons.

A striking observation from our analysis of pseudobulk DEGs reveals that around 15% of these DEGs overlap with SFARI gene candidates and display significant dysregulation in deep layer excitatory neurons, particularly the L5-6 ExN_1, L5-6 ExN_2, and Im L6 ExN clusters (**Supplemental Figure 1**). Interestingly, expression levels of these genes were elevated in KOs compared to WTs, hinting at the possible loss of a repressive mechanism. Despite the majority of these genes not being identified as direct targets in the E14 MYT1L CUT&RUN analysis, it is important to note that the CUT&RUN dataset only includes gene targets based on MYT1L occupancy in promoter regions, omitting potential targets influenced by distal regulatory elements, as it remains challenging to systematically link long-distance enhancers to specific gene targets. Nevertheless, the observed differential expression allows us to hypothesize that MYT1L may function as a transcriptional regulator, influencing SFARI gene expression directly or indirectly, or through mechanisms like epigenetic memory. Ultimately, the disrupted pathways we’ve identified represent a core set of pathways that are critical for proper neurodevelopment.

Overall, our comprehensive analyses across developmental stages and MYT1L deficiency’s impact underscore its pivotal role in neuronal maturation and development. These findings reveal that the developmental trajectory and transcriptional landscape of excitatory neurons are markedly altered by MYT1L

deficiency, with effects persisting from early neurogenesis through adolescence. This study not only advances our understanding of the genetic and molecular foundations of neuronal development, but also demonstrates how we can deeply characterize genetic perturbations at scale to investigate the enduring impact of MYT1L on the maturation and function of neurons.

## Methods

### Animals and tissue collection

All animal studies were approved by and performed in accordance with the guidelines of the Animal Care and Use Committee of Washington University in Saint Louis, School of Medicine and conform to NIH guidelines of the care and use of laboratory animals. The animals were housed in controlled environments with a 12-hour light-dark cycle, constant temperature and relative humidity, and *ad libitum* access to food and water. The C57BL/6-*Myt1l^em1Jdd^*/J (Myt1l S710fsX^12^; Jackson Laboratories 036428) line was maintained with breeding pairs consisting of a Myt1l Het and an in-house C57BL/6J mouse. The transgenic line was refreshed every 8-10 generations by backcrossing to freshly obtained C57BL/6J males and females from Jackson Laboratories. Upon weaning at P21, the animals were group-housed by sex and genotype. To obtain homozygous animals for embryonic studies, timed pregnant Myt1l Het x Het breeding pairs were set up and vaginal plugs were checked the following morning. The first day after the plug was found was considered to be E0.5. E14-14.5 embryos were rapidly dissected from the uteri of the mice in HBSS on ice. The pups were decapitated, and the brains were quickly extracted from the skulls. The meninges was removed, the forebrain was dissected, flash frozen in liquid nitrogen, and stored at -80°C. Tail tissue was collected from each embryo for gDNA isolation and genotyping. Cortical tissue from P21 pups were harvested and stored using the same method.

### Genotyping

Tissue (tail biopsy, ear punch, or toe clipping) was obtained from each animal and placed in a PCR tube. 100ul lysis buffer (25mM NaOH, 0.2mM EDTA, pH 12) was added to each tube and incubated at 99°C for 60 min in a thermocycler. Once the samples cooled to room temperature, 100ul 40mM Tris-HCl pH 5 was added to neutralize the alkaline lysis buffer. The crude lysate containing genomic DNA (gDNA) was stored at 4°C. Three reactions were performed for each animal to genotype the WT allele, MYT1L mutant allele, and SRY to determine sex (see **Table 1** for sequences). The PCR conditions for genotyping with allele specific PCR primer pairs involved mixing 1ul of the crude gDNA with 5ul Phusion High-Fidelity PCR Master Mix, 1ul 10uM MYT1L F/R primer mix, 1ul 10uM B-actin F/R primer mix, and 2ul ddH2O. Thermocycling conditions were as follows: 98C for 3 min; 35 cycles of: 98C for 10 sec, 61C for 20 sec, 72C for 20 sec; 72C for 5 min; and 4C hold. For SRY, 1ul crude gDNA was added to a master mix containing 5ul OneTaq Quick-Load 2X Master Mix, 1ul 10uM SRY primer mix, 1ul 10uM B-actin primer mix, and 2ul ddH2O. Thermocycling conditions were as follows: 94C for 3 min; 35 cycles of: 94C for 10 sec, 60C for 20 sec, 68C for 20 sec; 68C for 5 min; and 4C hold. Multiplexing B-actin not only confirms the presence of gDNA, but also minimizes non-specific amplification of the MYT1L mutant band in WT samples. PCR products were run on a 1% agarose gel and visualized with GelRed.

### Injection of AAV-Calling Cards reagents

Calling Cards is a method to longitudinally record protein-DNA interactions over time in tissues^40–42^. The constructs hyPB and H2b-tdT-SRT (Addgene 203393) were packaged into AAV9 viral particles by the Hope Center Viral Vectors Core at Washington University. The titer was determined by qPCR and standardized to 1x10^13^ vg/ml. A step-by-step protocol for transcranial injections is described in Yen et al^42^. Briefly, the AAVs were mixed 1:1 and transcranially injected into the ventricles of P0-1 pups from MYT1L WT x Het breeding pairs. At P7, toe tissue was collected for genotyping. At P21, the animals were deeply anesthetized with isoflurane and perfused with ice cold DPBS. The brain was harvested, tdTomato fluorescence was verified using a handheld fluorescence flashlight (Nightsea Xite-GR), the cortex was dissected, and the tissue was flash frozen in liquid nitrogen and stored at -80C. The tissue was processed following the “nuclei isolation and fixation” section below.

### Nuclei isolation and fixation

In this study, nuclei from E14 and P21 prefrontal cortices were isolated from flash frozen tissue. The brain tissues were Dounce homogenized in ice-cold homogenization buffer (10mM Tris-HCl pH 7.4, 10mM NaCl, 3mM MgCl2, 1mM DTT, 1X cOmplete EDTA-free Protease Inhibitor (Roche 4693132001), and 0.2U/ul RNasin Inhibitor (Promega N2515) using a 2ml KIMBLE KONTES Dounce Tissue Grinder (DWK 885300-002) with 15 strokes with the “A” large clearance pestle, followed by 15 strokes of the “B” small clearance pestle. The homogenate was transferred to a 15 ml centrifuge tube. Walls of the homogenizer tubes were washed with 1ml of homogenization buffer and combined with the homogenate in the 15ml tube. The nuclei were pelleted by centrifugation in a swinging bucket rotor at 500x g for 5 mins at 4C. The supernatant was aspirated and discarded.

For E14 samples, the pellets were washed twice with 1ml nuclei wash buffer (DPBS, 1% BSA, and 0.2U/ul RNase inhibitor), filtered through a 40um Flowmi cell strainer (Millipore Sigma BAH136800040), and counted using a hemocytometer with Trypan Blue.

For P21 samples, the pellets after the first centrifugation were resuspended in 1ml homogenization buffer. A gradient centrifugation step using 25:35 Iodixanol was performed to purify the nuclei from cellular debris and myelin generated during tissue dissociation. To make the 25% Iodixanol layer, 1ml 50% iodixanol was added to 1ml of the homogenate containing the nuclei and debris. This was carefully layered on top of 2ml 35% iodixanol in a clear polycarbonate tube (Beckman 355672) and centrifuged at 10,000x g for 30 min at 4C with deceleration turned off. After the gradient centrifugation, myelin and cellular debris remaining at the top of the 25% iodixanol layer and was aspirated and discarded. The purified nuclei at the interface of the two Iodixanol layers was collected and transferred to a clean 15ml centrifuge tube. The volume was brought up to 6 ml with nuclei wash and resuspension buffer and pelleted by centrifuging at 500x g for 5 min at 4C. The supernatant was carefully removed, washed once with nuclei wash buffer to ensure removal of carryover Ioxidanol, filtered through a 40um Flowmi cell strainer, and counted using a hemocytometer with Trypan Blue.

For both E14 and P21 samples, 500k-2.5M nuclei were resuspended in 500ul calcium and magnesium-free DPBS and used as input into the ScaleBio Sample Fixation Kit (Scale Biosciences 2020001) protocol according to manufacturer’s instructions. After fixation, the nuclei were counted once more and checked for quality using a microscope with a 60x objective. The nuclei were then stored at -80C until all samples have been collected and fixed.

### Single-nucleus RNAseq library preparation

Libraries were prepared from fixed E14 and P21 nuclei separately. For the E14 timepoint, a total of 9 samples (3 biological replicates of a mix of males and females per MYT1L WT, Het, and KO genotypes) were used. For the P21 timepoint, a total of 12 samples consisting of 3 biological replicates per sex per MYT1L WT and Het genotypes were used. The day of the library preparation, the frozen fixed nuclei were thawed on ice and each sample was counted twice using a hemocytometer.

For the E14 samples, the ScaleBio Single Cell RNA Sequencing Kit v1.0 (Scale Biosciences 2020008) was used according to manufacturer’s instructions. Nuclei from each sample were loaded at 10,000 nuclei per well to the 96-well Indexed RT Oligo Plate to add the RT barcode and UMI onto each transcript during reverse transcription. By loading each sample into a distinct set of wells, the RT barcodes can serve as sample identifiers, enabling all genotypes to be processed on the same plate in a single batch per age. The nuclei from each well were then collected and pooled using the Scale Biosciences’ supplied collection funnel, mixed, and distributed across the 384-well Indexed Ligation Oligo Plate where the Ligation Barcode was added to each UMI-RT barcoded transcript. Then, the nuclei were once again collected and pooled using another collection funnel and counted with a hemocytometer with Trypan Blue. A total of 1,600 nuclei were distributed per well of the 96-well Final Distribution Plate. In each well, second strand synthesis was performed followed by a cleanup step. The PCR products were then tagmented followed by an indexing PCR step to add a third barcode to each well. 5ul from each of the 96 libraries were pooled and cleaned using 0.8X SPRIselect beads (Beckman Coulter B23317). The average fragment size of the final library was quantified using a High Sensitivity D5000 Screentape (Agilent). The library concentration was quantified using the NEBNext Library Quant Kit for Illumina (New England Biolands E7630S). The libraries were sequenced on a shared S4 flowcell on a NovaSeq6000 (Illumina) instrument to a target depth of 10,000 reads per nucleus.

For the P21 samples, the protocol described above was followed through the cleanup step. Prior to tagmentation, 3ul (half of the total volume) was transferred to a clean 96-well PCR plate to create a Calling Cards Final Distribution Plate. This plate was set aside to pilot single-nucleus Calling Cards (snCC) library preparation, the results of which will be reported in a future methods paper. The remaining 3ul was used for the remainder of the ScaleBio protocol with slight modifications. To account for the reduced volume of template input, the volumes for all subsequent steps have been halved to keep all reaction proportions the same. Additionally, the Indexing PCR program was increased to 16 cycles instead of 14. The libraries were pooled, cleaned, and quantified as described above according to manufacturer’s instructions. This library pool was sequenced on a shared 25B flowcell on a Novaseq X (Illumina) instrument to a target depth of 10,000 reads per nucleus.

### snRNAseq analysis

Base calls were converted to fastq format and demultiplexed by Index1 barcode by the Genome Technology Access Center at the McDonnell Genome Institute (GTAC@MGI). Combinatorial barcode demultiplexing, barcode processing, adapter trimming, read mapping to the mm10 reference genome, single-nuclei counting, and generation of the feature-barcode matrices were done using ScaleBio Single-cell RNA Nextflow Workflow v1.4 (https://github.com/ScaleBio/ScaleRna). The count matrices were brought into Seurat for downstream analyses. For quality control, a UMI-gene cutoff of 800-6000 UMIs and 300-3000 genes was used, followed by filtering out multiplets by DoubletFinder^43^ for each sample. After filtering, a total of 216,830 nuclei across all samples remained, with a median of 3,204 UMIs and 1,819 genes per nucleus. The count matrices were log2 normalized, centered, and scaled using a scaling factor of 10,000. The top 3000 most variable genes were identified using dispersion and mean expression thresholds. PCA was then performed followed by dimensionality reduction by UMAP and unsupervised clustering using the Louvain algorithm using the FindClusters function of Seurat. A range of values was tested for the resolution parameter and a clustering tree was plotted using clustree^44^ to determine 0.8 as the optimal resolution for the E14 dataset. Cluster marker genes were defined by grouping the clusters by genotype, setting logfc.threshold=0.5 and min.pct=0.25, and comparing the fold changes between pct.1 and pct.2 using FindConservedMarkers. Cell type annotations were done referencing literature and the ABC atlas. Pseudobulk differential expression analysis between genotypes within cell types was performed using DESeq2^28^. Pseudotime and trajectory analysis was done using Monocle3^34^. To prepare the data for RNA velocity analysis, .loom files containing the spliced and unspliced counts matrices were constructed from the .bam files and feature-barcode matrices using velocyto^45^ and the mm10 genome. RNA velocity was then computed with the dynamical model of scVelo^46^, which was then used in CellRank’s^47^ VelocityKernel to compute a macrostate transition matrix to classify initial, terminal or intermediate cell states.

For analysis of the P21 brains, a similar workflow as described above was used. After quality control, 96,505 nuclei remained with a median of 3,447 UMIs and 1,597 genes per nucleus. Cell type annotations was done exclusively using the ABC Atlas^19^. The total library counts were normalized, centered, and scaled using a scaling factor of 10,000. The dimensionality of the data was reduced first with principal component analysis on 100 components based on the top 3,000 most variable genes. The graph was then embedded and visualized in two dimensions using UMAP. The nuclei were clustered using the Louvain algorithm using the FindClusters function of Seurat. A range of values was tested for the resolution parameter and a clustering tree was plotted using clustree to determine 0.8 as the optimal resolution for the P21 dataset.

### Statistics

No statistical methods were used to predetermine sample sizes. Samples were generally littermates and genotypes were assigned randomly by the sperm at conception, with no input from investigators. The investigators were not blinded to the samples, but all samples were processed in parallel in the same batch.

### Data Availability

The data generated in this study can be downloaded in raw and processed forms from the Gene Expression Omnibus (pending) and Neuroscience Multi-omic Data Archive (NeMO). The E14 MYT1L CUT&RUN dataset was downloaded from Gene Expression Omnibus (GEO: GSE222072).

### Code Availability

The ScaleRna Nextflow pipeline for processing raw Fastq reads to feature-barcode matrices is available at Github (https://github.com/ScaleBio/ScaleRna). The code needed to reproduce the key findings of this paper is found at Bitbucket (pending).

## Acknowledgements

We thank members of the Dougherty and Mitra laboratories for helpful discussions and feedback, particularly M. Vasek for critical reading and copyediting of the manuscript; Y. Li for helpful analysis suggestions; the DNA Sequencing and Innovation Lab (DSIL) and the Genome Technology Access Center at the McDonnell Genome Institute (GTAC@MGI) for their sequencing expertise and services; and F. Schlesinger and B. Biddy for bioinformatic support for the ScaleBio pipeline. This work was funded by grants from the National Institute of Mental Health (RF1MH117070, RF1MH126723, and R01MH124808 to J.D.D. and R.D.M.). A.Y. was supported in part by T32HG000045 from the National Human Genome Research Institute.

## Author contributions

Project conceptualization: A.Y., R.D.M., and J.D.D. Method development, experiments, and data collection: A.Y., X.C., D.S., F.L., M.C., J.H-L, and J.C. Formal analysis: A.Y., Y.W., R.D.M., and J.D.D. Figures and data visualization: A.Y., R.D.M., and J.D.D. Writing-original draft: A.Y. and J.D.D. Writing-review and editing: A.Y., J.C., R.D.M., and J.D.D. Project coordination: A.Y., R.D.M., and J.D.D. Funding acquisition: R.D.M. and J.D.D.

## Supplemental Figures

**Supplemental Figure 1.**
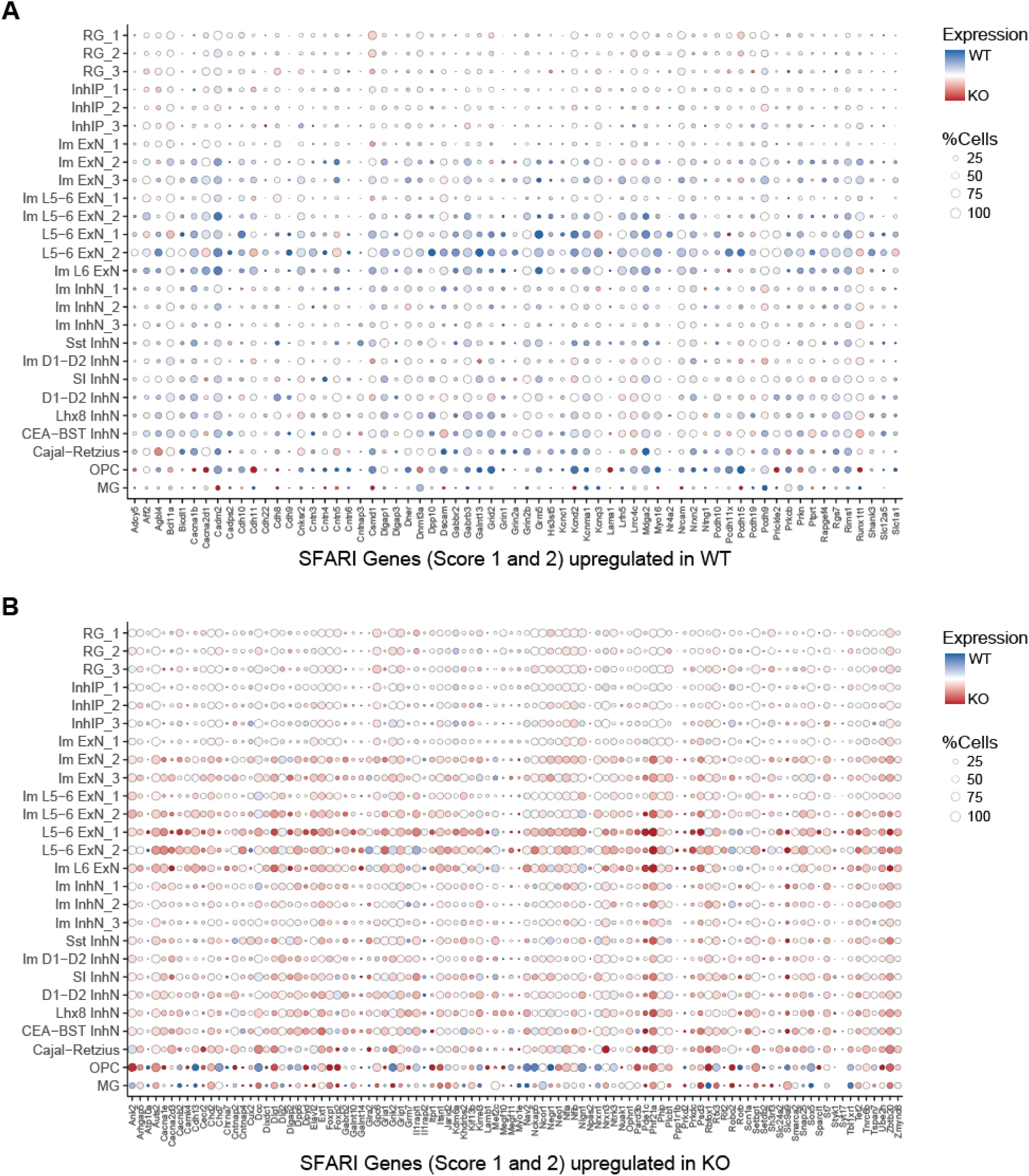
Cell type-specific dysregulated genes associated with autism. 164 differentially expressed genes from WT and KO E14 were found to be high confidence SFARI genes with a score of 1 or 2. The dotplot shows the genes as columns and cell types as rows. The color scale represents if the gene is upregulated in WT or KO samples, and the size of each circle represents the percentage of cells expressing the gene. This figure demonstrates an increased disruption of the SFARI genes in deep layer neurons (L5-6 ExN_1, L5-6_ExN_2, and Im L6 ExN clusters) and a general trend for these SFARI genes to be upregulated in KO samples.

**Supplemental Figure 2.**
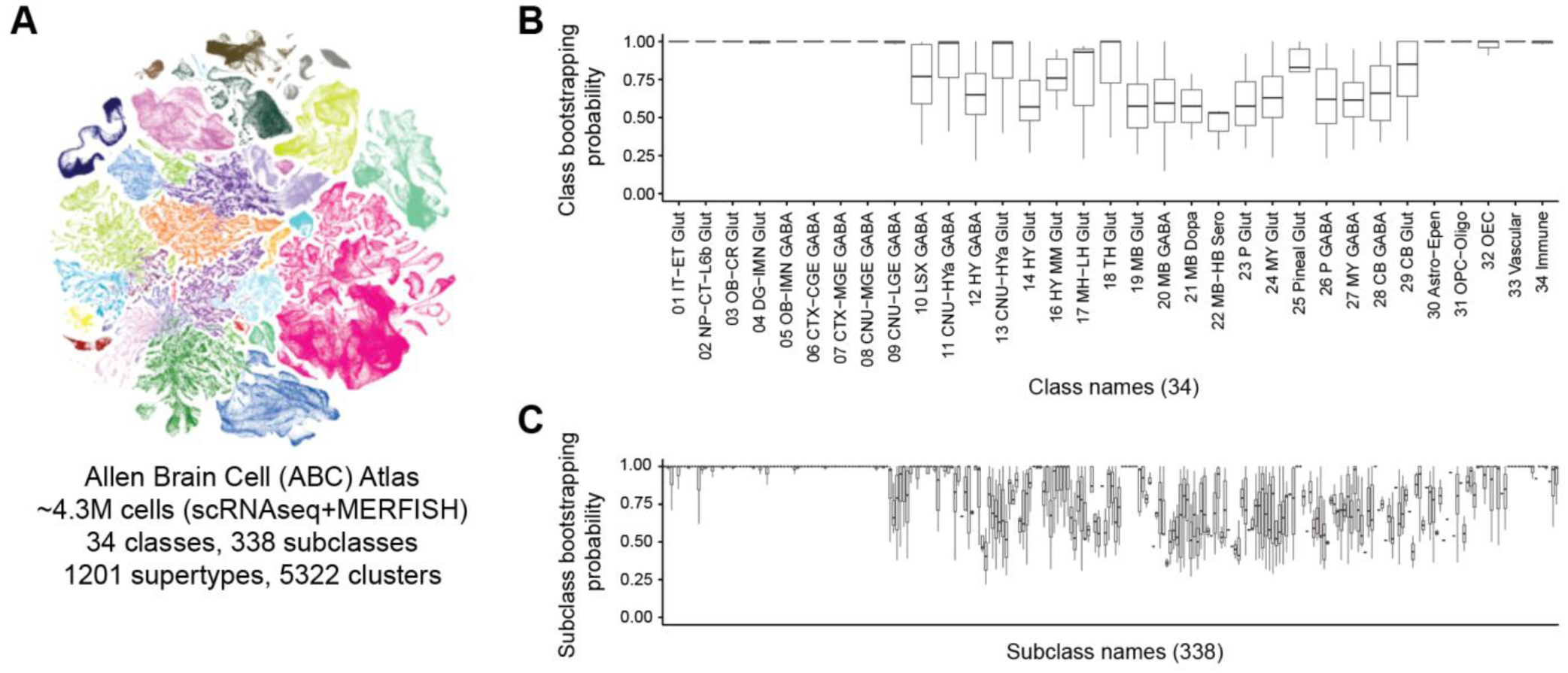
Standardized cell annotation of P21 dataset using the Allen Brain Cell Atlas. (A) Representative UMAP from the Allen Institute’s Allen Brain Cell (ABC) Atlas (https://portal.brain-map.org/atlases-and-data/bkp/abc-atlas) that consists of transcriptomes and anatomical location of millions of cells. Using MapMyCells (https://portal.brain-map.org/atlases-and-data/bkp/mapmycells) and hierarchical correlation mapping, cells from this study were mapped onto the atlas. For each cell, a random set of 90% of the marker genes was selected, then mapped to the atlas by traversing the taxonomy by starting with classes, then proceeding to subclasses, supertypes, and clusters. This was repeated 100 times to obtain the bootstrapping probability. The bootstrapping probability for class is shown in **(B)** and subclass is shown in **(C)**. Labels with a high bootstrapping probability (close to 1) are considered high confidence labels and subclass names were used to annotate nuclei from this study.

**Supplemental Figure 3.**
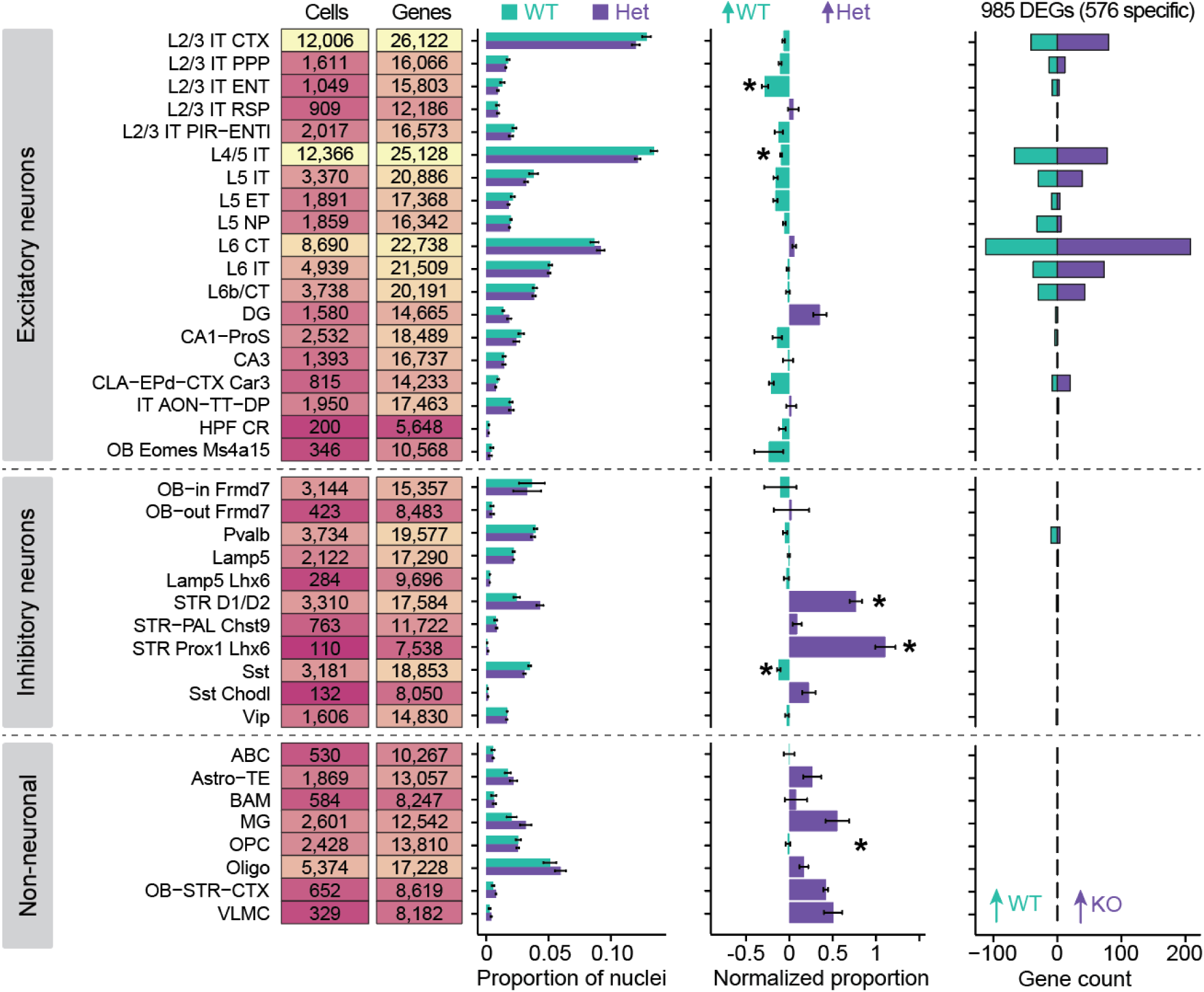
Single nucleus transcriptional profiling of P21 cortex in MYT1L animals. Summary plot showing the numbers of nuclei and genes detected in all clusters from P21 animals. The excitatory neuron subset was shown in Figure 4B. Bar plots show the average relative proportions of nuclei in each annotated excitatory neuron cluster for MYT1L WT and Het genotypes. These proportions are normalized to WT (center bar plot). The right bar plot shows the number of pseudobulk differentially expressed genes that are upregulated in WT (cyan) or upregulated in Het (purple).

